# Integrated approaches to identifying cryptic bat species in areas of high endemism: the case of *Rhinolophus andamanensis* in the Andaman Islands

**DOI:** 10.1101/561860

**Authors:** Chelmala Srinivasulu, Aditya Srinivasulu, Bhargavi Srinivasulu, Gareth Jones

**Affiliations:** Natural History Museum and Wildlife Biology & Taxonomy Lab, Department of Zoology, University College of Science, Osmania University, Hyderabad, 500007, Telangana, India; Systematics, Ecology & Conservation Laboratory, Zoo Outreach Organisation (ZOO), No 12, Thiruvannamalai Nagar, Saravanampatti-Kalapatti Road, Saravanampatti, Coimbatore, 641035, Tamil Nadu, India; Biodiversity Research and Conservation Society, 303 Nestcon Orchid, Kanajiguda, Tirumalgiri, Secunderabad, 500015, Telangana, India; School of Biological Sciences, Life Sciences Building, 24 Tyndall Avenue, Bristol BS8 1TQ, UK

## Abstract

The diversity of bats worldwide includes large numbers of cryptic species, partly because divergence in acoustic traits such as echolocation calls are under stronger selection than differences in visual appearance in these nocturnal mammals. Island faunas often contain disproportionate numbers of endemic species, and hence we might expect cryptic, endemic species to be discovered relatively frequently in bats inhabiting islands. Species are best defined when multiple lines of evidence supports their diagnosis. Here we use morphometric, acoustic, and molecular phylogenetic data to show that a horseshoe bat in the Andaman Islands is distinct in all three aspects, supporting its description as a newly described endemic species. We recommend investigation into possible new and endemic bat species on islands by using integrated approaches that provide independent lines of evidence for taxonomic distinctiveness. We provide a formal description of the new species – *Rhinolophus andamanensis* Dobson, 1872.

## Introduction

Cryptic species represent an important and long-neglected component of biodiversity, and cryptic taxa often fill distinct ecological niches that merit specific conservation challenges [1]. The number of bat species described has increased dramatically in recent years: current estimates recognise over 1300 bat species [2], an increase of more than 40% since 1993 [3]. This increase has been partly due to a recent surge in research on bats, and also because many cryptic species have been discovered by modern integrative techniques including echolocation call analyses [4,5] and molecular phylogenetics [6,7].

The basis for recognising new cryptic species is challenging and is strongest if multiple and independent lines of evidence are used [8]. Reproductive isolation may occur through post-mating barriers and differences in genital morphology. In bats, the baculum (os penis) is often different in cryptic taxa, and this may lead to mechanical incompatibility during mating [9], and reproductive isolation. Hence taxa that differ in echolocation calls, gene sequences, and bacular morphology are strong candidates for being distinct species, even if their overall morphology and appearance may be superficially similar to other taxa. In echolocating bats, acoustic traits such as echolocation calls may be selected to diverge more readily than traits associated with visual appearance, as the sensory world of these bats is dominated by sound [10].

Although cryptic bat species have been described in regions where research has been long-established (e.g. the discovery that pipistrelles in Europe comprised two widespread and abundant taxa that echolocate using different peak call frequencies [7,11]), the potential for the discovery of cryptic taxa may be greatest in regions where little research has been conducted. DNA barcoding studies suggest that the number of bat species in Southeast Asia may be double the number currently described [12]. Cryptic species may be especially prevalent in areas of high endemism and in biodiversity hotspots, especially if the potential for speciation in such regions is considerable.

The Andaman Islands form an archipelago in the Bay of Bengal, which host many endemic species of animals and plants, including mammals [13]. They are included in the Indo-Burma biodiversity hotspot [14] and comprise 200 – 300 islands and islets [15] which provide considerable potential for allopatric speciation. The islands are >1200 km from mainland southeast Asia and were formed when the Indian tectonic plate collided with the Burma Minor plate about 4 million years ago [16,17]. Their geographical isolation from the mainland over long time periods provides considerable potential for the evolution of endemic taxa, especially for taxa with low aspect ratios and wing loadings (and hence limited dispersal abilities), such as horseshoe bats.

In this paper, we test the hypothesis that the intermediate horseshoe bat currently described as a subspecies of *Rhinolophus affinis* (*R. affinis andamensis*) from the Andaman Islands is distinct in terms of echolocation calls, bacular morphology, and mtDNA sequences in comparison with mainland forms. We use these multiple lines of evidence to propose the elevation of the subspecies to specific status.

Dobson [18] proposed that *Rhinolophus andamanensis* n.sp. resembled *Rhinolophus affinis*, and suggested that the taxon “may be referred to the same section of the genus [as *R. affinis*]”. Despite listing *R. andamanensis* as a distinct taxon, Ellerman and Morrison-Scott [19] suggested that it may represent *R. affinis*. Y.P. Sinha [20] compared *R. andamanensis* and *R. affinis superans* and found that *andamanensis* showed more affinities with *superans* than with the nominate forms, and also considered *andamanensis* as a subspecies of *R. affinis*. His claim was based on an erroneous interpretation of the earlier work by Dobson [18] and Ellerman and Morrison-Scott [19]. While Corbet & Hill [21] and Bates & Harrison [22] synonymised *R. andamanensis* under *R. affinis* without clear justification, it was listed as a subspecies of *R. affinis* in Simmons [23] and Srinivasulu & Srinivasulu [24]. After the initial collection in 1872, this taxon was subsequently collected from Interview Island in 1959 [20], Diglipur in 1980 [25], and Paget Island, East Island, Chalis Ek, Saddle Peak in North Andaman, Interview Island, Baratang Island, and Little Andaman in 2002 [26].

Our earlier analyses suggested that the taxon *andamanesis* is distinct from *R. affinis*, indicating that further study on this species is needed to assess its taxonomic status [27]. The present study provides evidence based on statistical analysis of morphometrics, molecular phylogenetics of fresh specimens collected from various sites in the Andaman Islands between 2012 and 2016, and comparative studies of museum specimens housed in various collections. We use this information, supported by evidence from echolocation call analysis to confirm the specific status of this taxon. A detailed description of *R. andamanensis*, primarily based on the holotype and fresh specimens is also provided, as the original description provided by Dobson [18] is too short and lacks details on characters. We use our approach to show how data on the species’ morphology, echolocation call parameters, baculum characters, and phylogenetics can provide multiple independent lines of evidence to describe new cryptic bat species.

## Materials and Methods

### Legal and Ethical Statements

Fresh voucher specimens were collected under the Study-cum-Collection permit (No. CWLW/WL/134/235, dt. 13 October 2014) issued by the Andaman and Nicobar Forest Department, Government of Andaman and Nicobar Islands. Captured bats were handled in strict accordance with good animal practices and according to the American Society of Mammologists guidelines [28]. As the voucher specimens were collected, euthanized and preserved in field ethics approval from Institutional Animal Ethical Committee was not required for this study.

### Study specimens and study sites

Thirty-three specimens were studied including three specimens (including the holotype, ZSI Reg. No. 15561) from the National Collection at Zoological Survey of India (ZSI), Kolkata; three specimens from the Harrison Zoological Museum (HZM) at the Harrison Zoological Institute (HZM), Sevenoaks, UK (misidentified as *Rhinolophus yunanensis*); and 27 specimens collected during the present study [four specimens from Interview Island (12.991° N, 92.707° E), North Andaman; two specimens from Chipo (13.527° N, 93.013° E), North Andaman; eleven specimens from different caves on Baratang Island (12.090° N, 92.749° E), Middle Andaman; four specimens from Burmadera (12.860° N, 92.882° E), Middle Andaman; three specimens from Pathilevel (13.208° N, 93.009° E), Middle Andaman; two specimens from Ramnagar (13.077° N, 93.021° E), Middle Andaman; and one specimen from V.K. Pur (10.726° N, 92.576° E), Little Andaman].

### Morphometrics

External measurements on live specimens and craniodental measurements on the extracted skulls were taken using digital Vernier calipers (Mitutoyo make, to the nearest 0.01 mm). The following external and craniodental measurements were taken: External: FA, forearm length; E, ear length; Tl, tail length; Tib, tibia length; Hf, hindfoot length; 3mt, third metacarpal; 4mt, fourth metacarpal; 5mt, fifth metacarpal; 1ph3mt, first phalange of third metacarpal; 2ph3mt, second phalange of third metacarpal; 1ph4mt, first phalange of fourth metacarpal; 2ph4mt, second phalange of fourth metacarpal; Craniodental: GTL, greatest length of the skull; CBL, condylobasal length; CCL, condylocanine length; CM^3^, maxillary toothrow; C^1^– C^1^, anterior palatal width; M^3^–M^3^, posterior palatal width; ZB, zygomatic breadth; BB, braincase breadth; CM_3_, mandibular toothrow; M, mandible length. Bacula were extracted and stained following the standard method [29].

### Statistical analysis

Specimen data on different subspecies of *Rhinolophus affinis* was sourced from the work of Ith and colleagues [30] for comparison with the taxon *andamanensis*. The bats were classified into five groups: Java, Borneo, North Malaysia, and South Malaysia [30], and Andaman. *A priori* multicollinearity tests were conducted on the morphological variables in R [31], and a principal component analysis (PCA) was done on all the morphological characters in R [31].

### Echolocation call analysis

Echolocation calls of *R. andamanensis* were collected from different locations in Andaman using a Petterson D500X bat detector (Pettersson Elektronik, Uppsala: frequency range 5–190 kHz, 500 kHz sampling rate). The calls were visualized in BatSound software (Pettersson Elektronik, Uppsala: FFT size 1024, Hanning window), and only the pulses with good signal-to-noise ratio were selected for the analysis. For the analysis, frequency at maximum energy (FMAXE) and duration were taken into consideration.

### Molecular analysis

Wing punches from the specimens were taken and dry-preserved using silica gel. Genomic DNA was then extracted using DNEasy Blood and Tissue kit (QIAGEN). A PCR was conducted to amplify partial cytochrome c oxidase subunit 1 (COI) gene sequence using standard forward and reverse [VF1d (5’- TTCTCAACCAACCACAARGAYATYGG-3’) and VR1d (5’- TAGACTTCTGGGTGGCCRAARAAYCA-3’)] primers [32]. The PCR reaction was performed in a 25 μl reaction volume containing 2 μl of template DNA, 12.5 μl of 2X reaction buffer (0.05 U/μL Taq DNA polymerase, reaction buffer, 4 mM MgCl2, 0.4 mM of each dNTPs), 0.5 μl of each primer, and 9.5 μl nuclease free water. The amplified PCR products were sequenced using an ABI prism 3730 sequencer (Applied Biosystems, USA) and Big dye terminator sequencing kit (ABI Prism, USA), and a 614 base pairs length of 5 sequences of cytochrome c oxidase subunit 1 (COI) gene was generated.

A total of 17 COI sequences of *R. affinis* and two sequences of *R. lepidus* (the outgroup taxon) obtained from GenBank, along with five sequences of *R. andamanensis* (present study) were used for the phylogenetic analysis (S2 Table). The sequences were aligned using MUSCLE [33] incorporated in MEGA6 [34] using default parameters. The best-fit maximum likelihood nucleotide substitution model was chosen in JModelTest2 [35,36], using the Bayesian Information Criterion (BIC) value for each model. The model used was the Hasegawa-Kishino-Yano + invariant sites (HKY+I, BIC = 3409.38) nucleotide substitution model [37].

To estimate the time to the most recent common ancestor (TMRCA) for *Rhinolophus andamanensis* and *R. affinis*, we used a mutation rate of 0.020 ± 0.005 per site per lineage per Myr [38]. A Bayesian inference of phylogeny was constructed using BEAST v1.8.2 [39] using a Yule speciation prior for a chain length of 50 million generations, sampling every 1000 generations. Two sequences of *Rhinolophus lepidus* were used to root the tree, and model performance was assessed by plotting likelihood scores against generations and checking the effective sample size (ESS) values in Tracer 1.6 [40]. The first 25% of the sampled trees were discarded as burn-in, and the final tree was created using TreeAnnotator v1.8.2 (part of the BEAST package) and visualised in FigTree v1.4.1 [41]. MEGA6 was used to calculate Kimura 2-parameter distances.

## Results

A total of 21 external and craniodental characters were used for conducting a PCA on 41 individuals of *Rhinolophus andamanensis* and *R. affinis* (Table 1). This generated five separate groups (Fig 1); four groups, representing *R. affinis* from Borneo, Java, Northern Malaysia, and Southern Malaysia overlapped with each other, and one group, representing *R. andamanensis*, was isolated from the rest. PC1 and PC2 explained 67.43% and 9.12% of the total variance respectively. In PC1, all loading scores were positive; in PC2, Tail length (TL) (positive, 1.60) and Forearm length (FA) (negative, −1.01) showed relatively large scores (Fig 1, S1 Table). These loadings fit with *R*. *andamanensis* having a larger body size than *R. affinis*, as confirmed by univariate measurements (S1 Table).

**Table 1.** Morphometric measurements (in mm) of Rhinolophus andamanensis and Rhinolophus affinis.

| *R. andamanensis* (n=27) |  |  |  | *R. affinis* (n=14) |  |  |
| --- | --- | --- | --- | --- | --- | --- |
|  | Mean±SD | Min | Max | Mean±SD | Min | Max |
| FA | 54.23±1.8 | 46.66 | 56.64 | 49.23±1.84 | 46.7 | 52.9 |
| TL | 24.3±2.86 | 17.92 | 30.97 | 21.53±2.1 | 18.8 | 25.8 |
| EL | 21.34±1.42 | 18.05 | 23.86 | 21.04±1.69 | 18.4 | 24.4 |
| TIB | 27.67±0.95 | 25.99 | 29.62 | 22.67±1.9 | 20 | 26.4 |
| HF | 12.56±0.99 | 9.74 | 14.66 | 10.21±0.78 | 9.04 | 11.6 |
| 3MT | 39.99±1.49 | 35.9 | 41.91 | 38.03±1.47 | 35.7 | 40.3 |
| 4MT | 41.79±1.56 | 37.47 | 43.84 | 38.83±1.55 | 36.5 | 41.8 |
| 5MT | 43.52±1.73 | 39.09 | 45.79 | 39.24±1.52 | 36.74 | 42.3 |
| 1P3MT | 16.34±0.68 | 14.58 | 17.34 | 14.53±0.76 | 13.3 | 16.2 |
| 2P3MT | 28.65±1.35 | 26.05 | 31.15 | 24.98±1.35 | 22.8 | 27.7 |
| 1P4MT | 11.1±0.68 | 9.47 | 12.44 | 9.93±0.84 | 8.2 | 11.5 |
| 2P4MT | 16.52±0.94 | 14.81 | 18.6 | 14.53±1.19 | 11.2 | 16.3 |
| GTL | 25.67±0.68 | 24 | 26.7 | 22.05±0.58 | 21.21 | 23.27 |
| CCL | 21.97±0.53 | 20.76 | 23.04 | 19.48±0.54 | 18.61 | 20.78 |
| **ZB** | 12.34±0.48 | 11.55 | 13.9 | 11.21±0.32 | 10.83 | 11.91 |
| **BB** | 10.9±0.69 | 9.38 | 11.75 | 9.58±0.28 | 9.27 | 10.14 |
| **CM<sup>3</sup>** | 9.85±0.30 | 8.98 | 10.32 | 8.79±0.27 | 8.41 | 9.38 |
| **CM<sub>3</sub>** | 10.82±1.36 | 10.06 | 17.51 | 9.18±0.31 | 8.7 | 9.82 |
| **M** | 16.79±1.58 | 9.68 | 17.84 | 15.24±0.43 | 14.58 | 16.07 |
| **M<sup>3</sup>-M<sup>3</sup>** | 8.68±1.11 | 5.68 | 9.68 | 8.12±0.35 | 7.65 | 8.86 |
| **C<sup>1</sup>-C<sup>1</sup>** | 7.11±1 | 5.77 | 9.59 | 5.61±0.25 | 5.12 | 6.11 |

**Fig 1.**
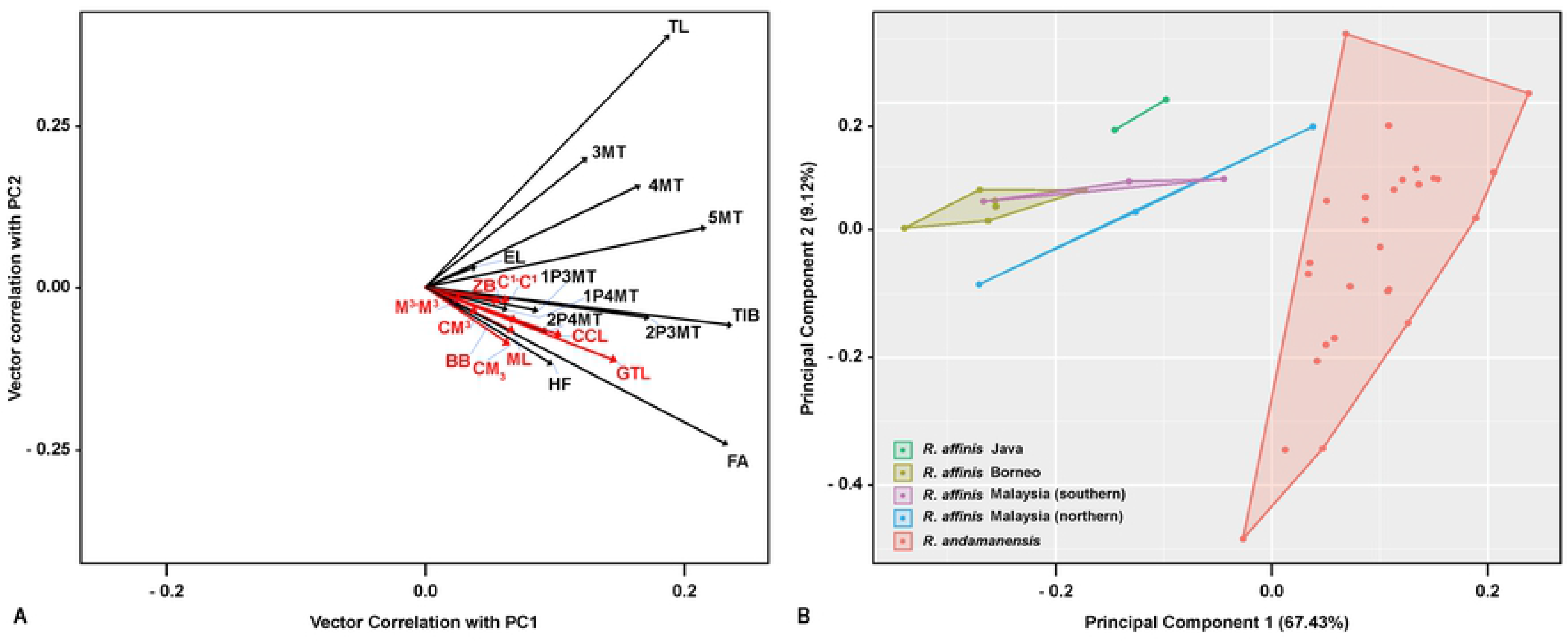
Principal component analysis of *Rhinolophus affinis* sensu lato and *Rhinolophus andamanensis* sensu stricto. A. vector loadings plot [cranial (red) and external (black) variables], B. PCA plot

### Echolocation calls

Echolocation calls of live specimens showed that the calls consist of a short upward broadband sweep followed by a long constant frequency component, followed by a short downward broadband sweep (Fig 2). The calls (n = 156) had a mean frequency of maximum energy of 57.65 ± 0.56 (SD) kHz (56.4 – 58.5 kHz) and a mean duration of 46.63 ± 8.59 ms (30 – 69 ms).

**Fig 2.**
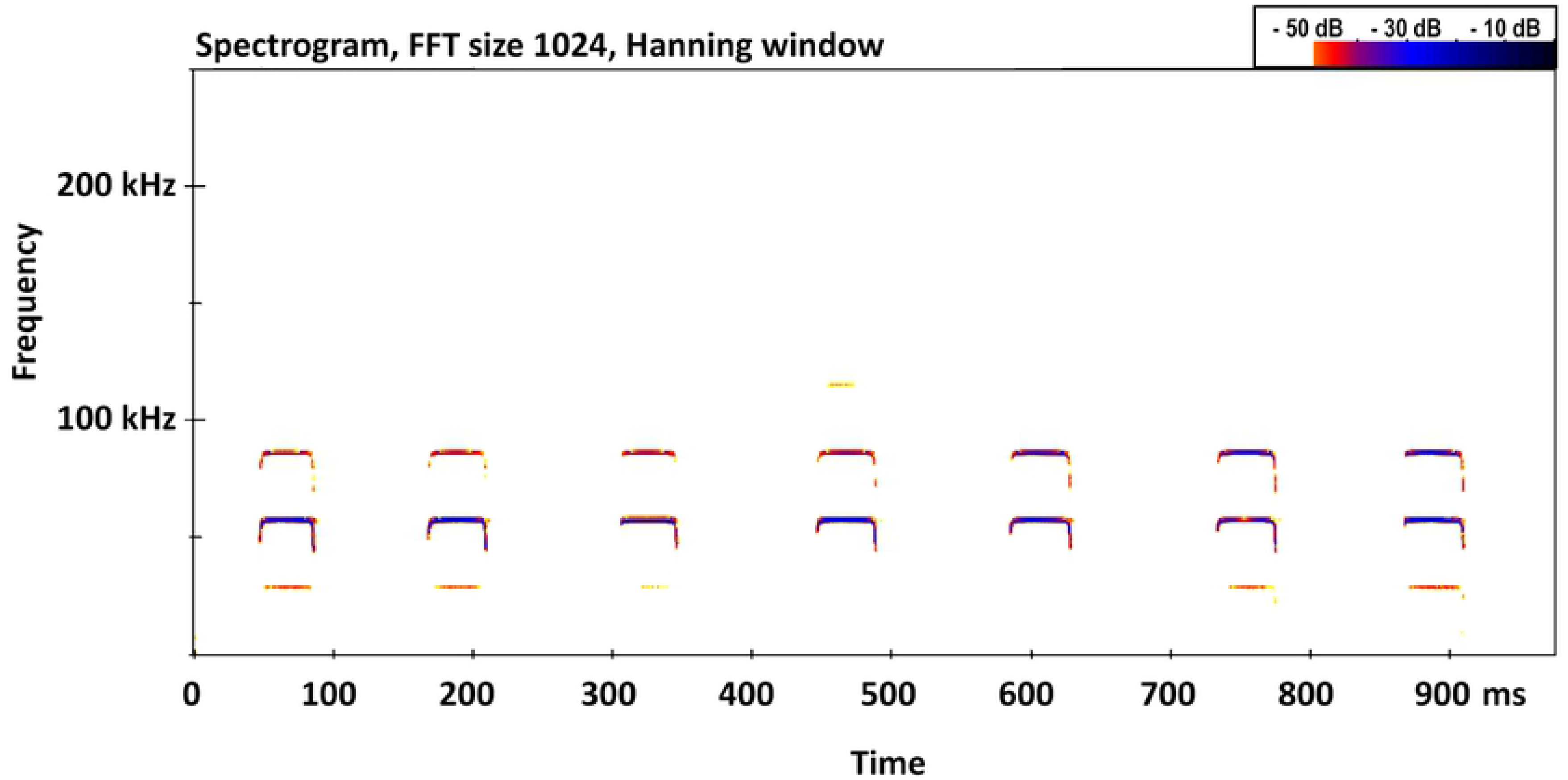
Spectrogram of echolocation calls of *Rhinolophus andamanensis* Dobson, 1872.

### Phylogenetic relationships

Phylogenetic analysis based on the selected COI gene sequences showed *Rhinolophus andamanensis* to be in a separate clade from all representatives of *R. affinis* (Fig 3). Within the *R. andamanensis* clade, two subclades were formed, the first includes individuals from Interview Island (NHM.OU.CHI.125.2014), Baratang Island (NHM.OU.CHI.31.2012), and Pathilevel (NHM.OU.CHI.113.2015), while the second includes individuals from V.K. Pur (NHM.OU.CHI.181.2015) and Chipo (NHM.OU.CHI.147.2015). The relationships between *R. andamanensis* and *R. affinis* are supported by high posterior probabilities. The K2P pairwise genetic distance between the *R. andamanensis* and *R. affinis* clades is 4.54%, 3.64–5.45% (mean, range); the highest mean distance was between *R. andamanensis* and individuals of *R. affinis* from Guangxi, Hunan, Vietnam, and Laos (4.68%), and the lowest mean distance was from *R. affinis* from Thailand (3.90%).

**Fig 3.**
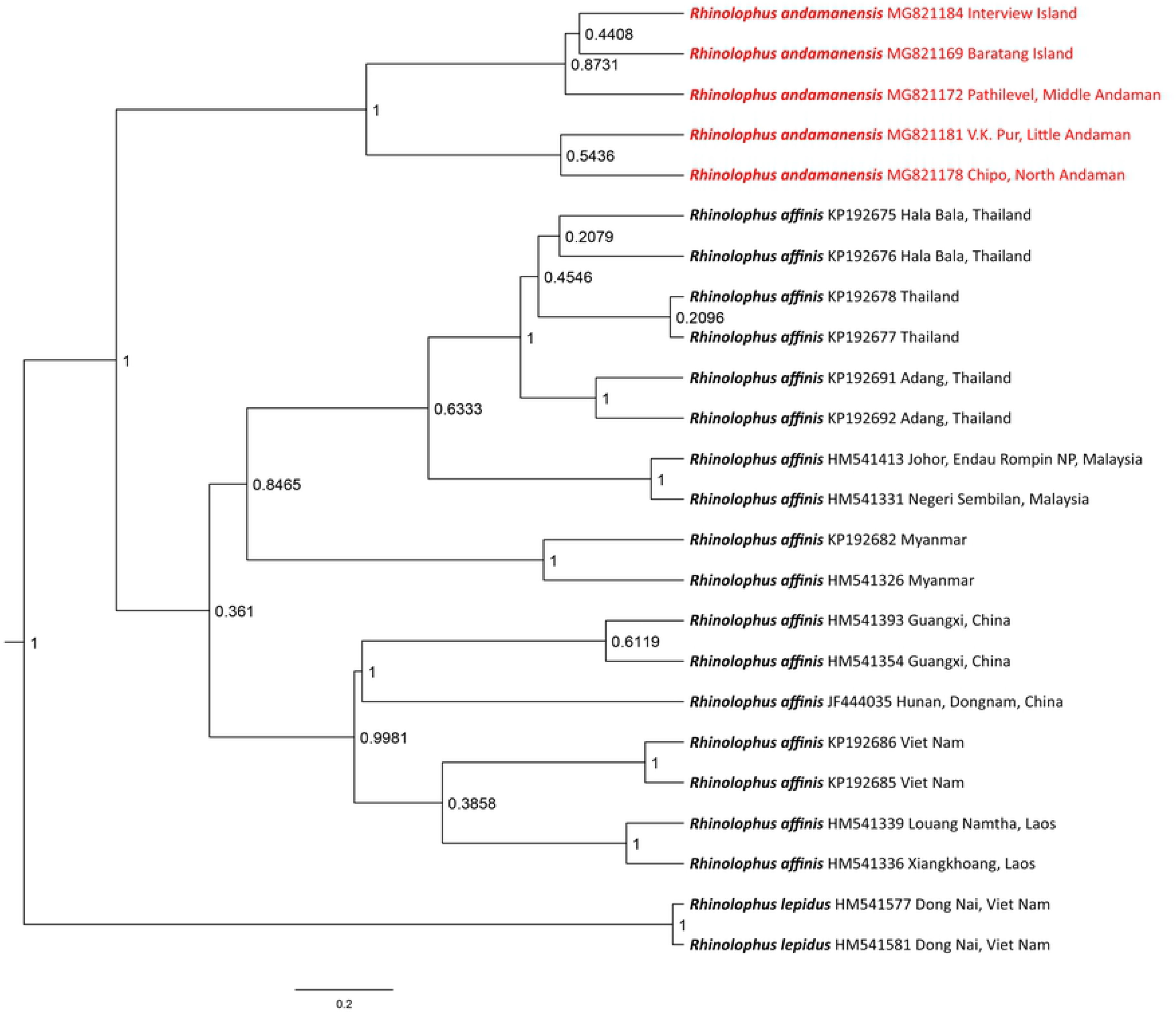
Maximum clade credibility topology estimated from Bayesian Inference (BI) of the Cytochrome C oxidase subunit I gene dataset using Hasegawa-Kishino-Yano + invariant sites (HKY+I, BIC = 3409.38). Values at the nodes are posterior probability values. *Rhinolophus lepidus* is used as an outgroup taxon.

Estimations of the time to the most recent common ancestor (TMRCA) were supported by high effective sample size values (ESS > 900) for all parameters. The inferred TMRCA for *R. andamanensis* vs. *R. affinis* was 1.39 My BP (95% CI: 0.46–2.76 My BP), which corresponds to the mid-Pleistocene epoch, and falls within a range of probable divergence times for the extant subspecies of *R. affinis* [28]. Sampling of additional markers is necessary to corroborate the intraspecific relationships among *R. affinis* populations.

### Systematic description

#### *Rhinolophus andamanensis* Dobson, 1872

Homfray’s Horseshoe Bat

##### Holotype

*Rhinolophus andamanensis* ZSI Reg. No. 15561, male, collected from South Andaman Island, Andaman Islands, India, 1872, by J. Homfray; specimen, and skull (damaged).

##### Diagnosis

A medium-sized bat with a forearm length ranging between 46.7–56.6mm. Ears tall and broad, with well-developed antitragus, tragus absent (Fig 4A). Horseshoe is broad and covers the whole of the muzzle, supplementary leaflet distinct; three mental grooves are observed on the lower lip (Fig 4B). Anterior border of the sella narrow above and wider below. Superior connecting process bluntly rounded off, inferior surface slightly bent, inferior extremity curved downward (Fig 4C). Sella roughly half the length of the lancet. Lancet narrow, triangular and tapered to a point; has three distinct cells (Fig 4B). Skull robust with a condylocanine length of 21.97±0.53mm. The maxillary toothrow (cm^3^) measures 9.85±0.30mm. pm^2^ is small but in the toothrow. pm_2_ is in the toothrow, and is half the height of pm_4_ and one third the height of the canine. Baculum narrow and long, base distinctly trilobed, the shaft curving gently when viewed laterally.

**Fig 4.**
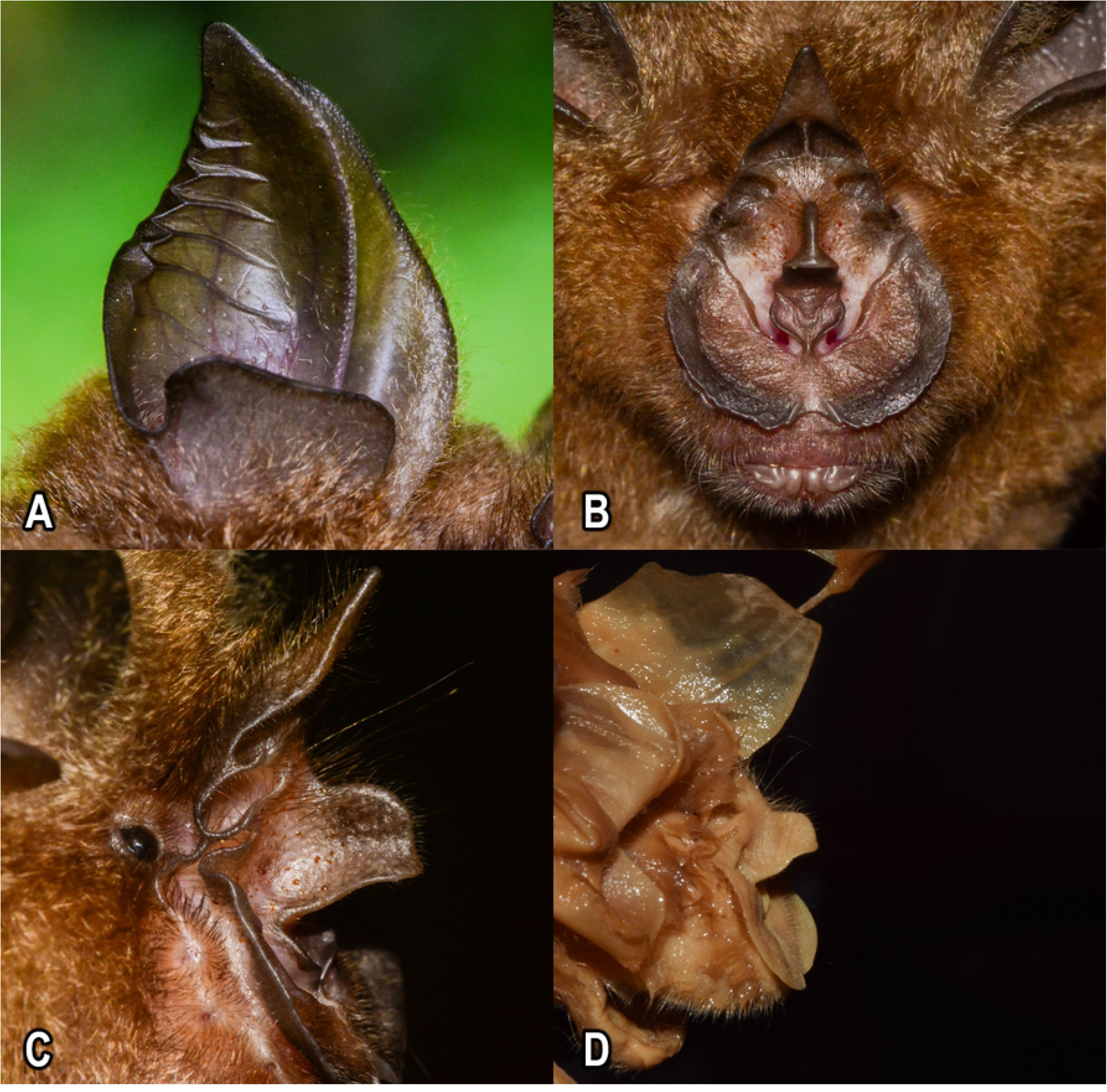
Live images (A-C) and holotype (ZSI Reg. No. 15561) (D) of *Rhinolophus andamanensis* Dobson, 1872. Ear pinna and antitragus (A), frontal view of the noseleaf and three mental grooves (B), and lateral view of the noseleaf showing shape of the sella (C) and the holotype (D).

##### External characters (based on holotype)

A medium-sized Rhinolophid with a forearm length of 52.98 mm. Ears tall (20.39 mm) and broad, ending with a pointed tip; antitragus well-developed, roughly triangular in shape, and half the height of the pinna; tragus absent. A slight concavity is observed on the outer border of the pinna just below the tip. Horseshoe broad and flat covering the whole of the muzzle; median emargination of horseshoe broad, well-developed and deep. Internarial cup of the horseshoe broad, taking up most of the narial space. The nares are located toward the base of the internarial cup, on either side. The internarial cup is attached to the horseshoe by means of a short stalk. Sella small, roughly half the length of the lancet; when viewed from the front, anterior border narrow above and wider below. Superior connecting process bluntly rounded off, the inferior surface is slightly bent, and the inferior extremity is curved downward (Fig 4D). Lancet is narrow, triangular and tapered to a point. Lancet has three distinct cells. Three mental grooves seen on the lower lip. Supplementary leaflet distinct. In the wing, the third metacarpal is 5.26% shorter than the fourth and 7.12% shorter than the fifth metacarpal. The first phalanx of the third metacarpal is 37.78% of the third metacarpal. The second phalanx of the third metacarpal is 69.81% of the third metacarpal. Wings and the interfemoral membrane attached to the tibia.

##### Colouration in preservation

Fur in preserved condition discoloured, with white hair bases and pale fawn tips.

##### Colouration in live condition

Fur colour varies from brown to bright orange. Ears and membranes dark in colour.

##### Craniodental characters (Fig 5)

Skull and the mandible of the holotype broken. The following description is based on skulls of recently collected specimens. Skull robust; greatest total length 24.0–26.7mm. Sagittal crest well-developed; extends up to the parietal region. Interorbital region broad. Rostrum bulged, with two well-developed nasal inflations. A bony septum divides the rostrum from the interorbital region; zygoma robust. Canines robust; the first upper premolar (pm^2^) is small and in the toothrow, between the second upper premolar (pm^4^) and the canine. The canine of the lower toothrow is tall. The second and fourth lower premolar (pm_2_ and pm_4_) are in contact as the third premolar (pm_3_) is small and extruded from the toothrow. The second premolar is one-third the height of the canine and half the height of the fourth premolar.

**Fig 5.**
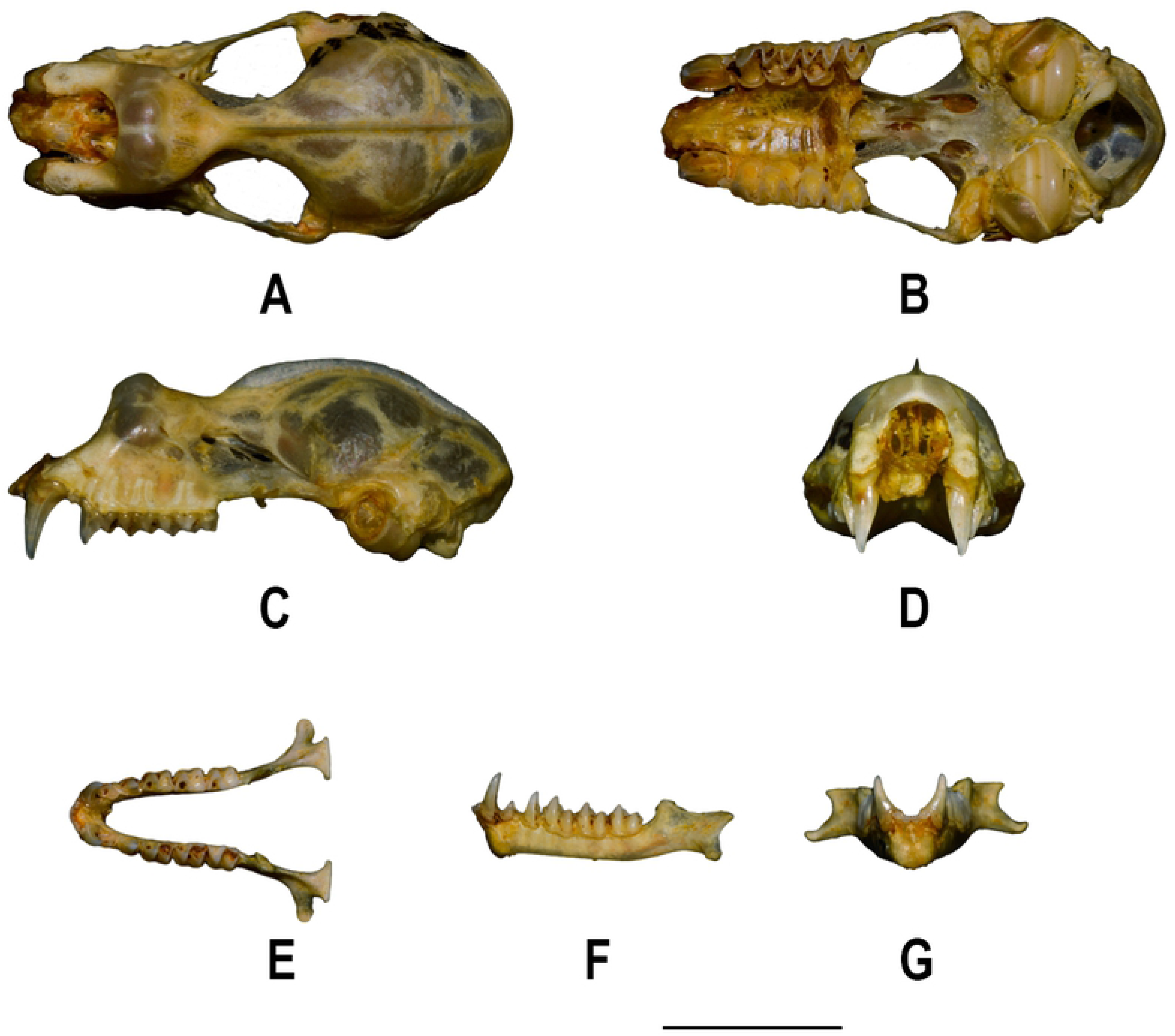
Skull and mandible of *Rhinolophus andamanensis* Dobson, 1872. Skull: A. Dorsal view, B. Ventral view, C. Lateral view, D. Frontal view; Mandible: E. Dorsal view, F. Lateral view, G. Frontal view (Scale: 5 mm).

##### Baculum

Baculum of holotype not extracted. Baculum in freshly collected specimens, 2-3 mm long with a slender shaft and a bulbous to a small base (Fig 6).

**Fig 6.**
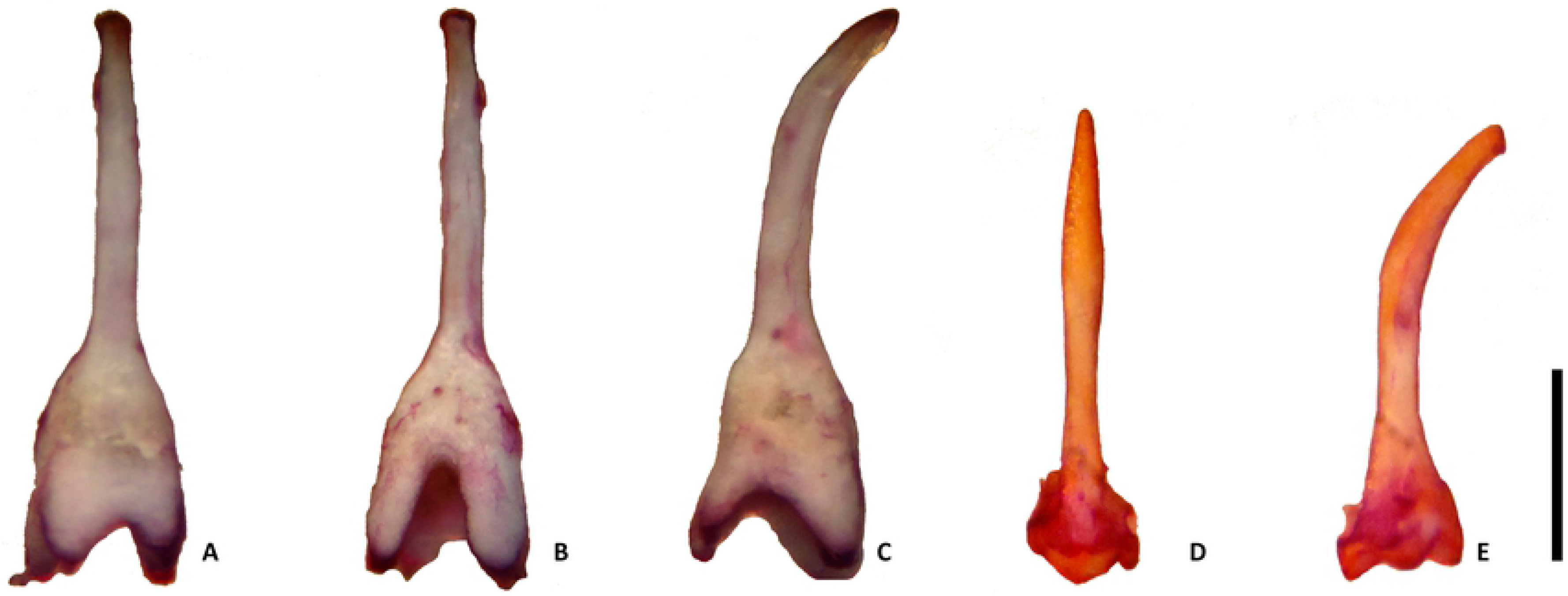
Baculum of *Rhinolophus andamanensis* Dobson, 1872. Typical form (A-C), and secondary type (D-E). A. & D. Dorsal view, B. Ventral view, C. & E. Lateral view (Scale: 1 mm).

##### Ecology

Little is known about the ecology of this species. Large colonies were found in limestone caves, forest caves, and sometimes in holes and hollows of large trees. It cohabits with *Rhinolophus cognatus*, *Hipposideros diadema masoni*, *H. gentilis*, *H. grandis*, and *Myotis horsfieldii*.

##### Distribution

*Rhinolophus andamanensis* is endemic to the Andaman Islands. It is distributed throughout the Andaman Archipelago - from north Andaman to Little Andaman.

## Discussion

We use morphological, acoustic, and genetic data to show that individuals hitherto referred to as *Rhinolophus affinis* from the Andaman Islands are distinct in all traits from mainland representatives of *R. affinis*, and deserve recognition as a new species. The long time and considerable distance of isolation from the mainland of Southeast Asia provided conditions suitable for allopatric speciation via morphological, acoustic, and genetic divergence from mainland ancestral forms. Our study shows that the taxon *andamanensis* is distinct from *R. affinis* under which it has previously been included as a subspecies, and is sufficiently divergent to be considered as a distinct species *Rhinolophus andamanensis* Dobson, 1872. Our TMRCA analysis suggests divergence from *R. affinis* about 1.4 My BP. Given that the islands began separation from the mainland about 4 My BP, our findings support the hypothesis that the new species evolved by allopatric speciation since the formation of the Andaman Islands archipelago.

Externally, *Rhinolophus andamanensis* is much larger than *R. affinis* (FA: 54.23±1.8mm vs. 49.23±1.84mm). In *R. andamanensis*, the skull is relatively robust (CCL: 21.97 ± 0.53mm vs. 19.7 ± 0.4 mm in *R. affinis*), and the palate is long (>50% of the length of the maxillary toothrow vs. <25% of the length of the maxillary toothrow in *R. affinis*). The bacular structure of the freshly collected specimens showed variations among individuals of *R. andamanensis*. Two major types of bacular structure were seen: the first type has a long shaft that curves ventrally, bulging slightly at the mid-section, and tapers to a pointed tip, the base is small and trilobed (Fig 6A-C), while in the second type the shaft is curved ventrally, is relatively thicker, and ends with a blunt tip. In the second type, from 25% up the height of the baculum, the shaft expands and continues to a four-pronged base with a deep V-shaped fissure, which appears low on the dorsal surface. Both the types of bacula were found among individuals collected from the same localities (Fig 6D-E). The bacular morphology of *R. affinis* studied in southeast Asia showed a divergent pattern between the Indochinese and the Sundaic subregions. The Indochinese bacula were smaller, had a curved shaft and a bulbous base, while the Sundaic population showed the presence of a larger bacula with a straighter shaft and a broad base [30].

The frequency of maximum energy (FMAXE) of the echolocation call of *R. andamanensis* ranged between 56.4–58.5 kHz [27]. In comparison, the echolocation calls of *R. affinis* sensu lato showed considerable variation across its distribution range [28]: calls of individuals from northern Thailand, Lao PDR, and Cambodia ranged from 70.0 to 79.9 kHz; northern populations in Vietnam called at 69.5–73.8 kHz, and southern populations at 81.2–84.5 kHz; populations south of the 7° latitude in Peninsular Thailand called at lower frequency (66.7–71.3 kHz), while those from Hala Bala, Narathiwat Province called at higher frequency (78 kHz). Hence the call frequency emitted by *R. andamanensis* is lower than that emitted by all populations of *R. affinis* studied, and the low call frequency is probably related to its larger body size, as call frequency scales negatively with body size in rhinolophid bats [42].

Phylogenetic analysis shows that *R. andamanensis* is distinct from *R. affinis*, under which it had been included due to similarities in morphology [19,20,22-24]. Although different populations of *R. andamanensis* branched in different subclades, the K2P pairwise genetic distance within the species clade was negligible (0–0.45%). Although *R. andamanensis* is principally a cave-dwelling species, it has also been observed roosting in tree hollows, and its echolocation calls have been recorded in the intervening areas of human habitation and forests, near streams, by the seashore, and in forests. This shows that this species is highly adaptable and is widely distributed throughout the island archipelago.

A recent haplotype network study [43] suggests that the populations of *R. andamanensis* from Little Andaman should be considered as distinct evolutionarily significant units. Our study, supported by K2P genetic distance, showed negligible genetic distance between individuals from Little Andaman and the rest of the population from mainland Andaman, suggesting that they have not yet diverged sufficiently to be considered distinct. Additionally, although individual variation in pelage colour, and shape and size of the baculum were evident, there exists little to no obvious structure in CO1 variation in *R. andamanensis* studied throughout the Andaman Islands to suggest that colour and baculum morphs may be isolated from one another.

Species richness on islands tends to decrease with increasing distance from the mainland [44], and islands often have relatively depauperate biota. Nevertheless, their isolation and the availability of vacant niches make islands (and especially archipelagos) ideal for allopatric speciation [45] and ultimately for the evolution of endemic taxa [46]. Because of powered flight, bats are often the only native mammals on oceanic islands, and many island bat taxa are evolutionarily and ecologically distinctive [47]. Island endemic bats are more threatened than non-island endemic species, and research on island endemics is lacking [48]. Because bat biodiversity comprises large numbers of cryptic species [12], we anticipate that large numbers of new cryptic bat species will be discovered on islands in the future, and recommend that their status is best validated by approaches that integrate morphology, molecular genetics, and - where appropriate - acoustics.

## Acknowledgements

We thank UGC-UKIERI Thematic Grants Programme, New Delhi for the funding for the work on the Andaman Islands. CS thanks DST-SERB, New Delhi, BS (UGC-PDF) thanks UGC, New Delhi, for research grants. We thank the Principal Chief Conservator of Forests (Wildlife) and Chief Wildlife Warden, Andaman and Nicobar Forest Department for study and collection permit, and the staff of the Andaman and Nicobar Forest Department for local logistics. We are thankful to Dr. Paul J.J. Bates and Dr. Malcolm Pearch for access to the collection at Harrison Zoological Institute, Sevenoaks, United Kingdom and helping us during our study. Mr. A. Gopi and Tauseef Hamid Dar for assistance in the field work, and the Head, Department of Zoology, Osmania University and UGC DSA-I (SAP II) for necessary facilities.

## Supporting information

S1 Table. Loading values of the first two principal components (PC1 and PC2) of the PCA analysis of external and craniodental morphometrics of *Rhinolophus andamanensis*.

S2 Table. Cytochrome C oxidase subunit 1 (COI) sequences of *Rhinolophus andamanensis* and *Rhinolophus affinis*, their collection localities, and GenBank accession numbers used for conducting the phylogenetic analysis; *Rhinolophus lepidus* was used as outgroup taxon.

